# Pituitary adenylate cyclase-activating polypeptide (PACAP) overexpression in the paraventricular nucleus of the thalamus alters motivated and affective behavior in female rats

**DOI:** 10.1101/2024.10.02.616278

**Authors:** Brody A. Carpenter, Breanne E. Pirino, Malcolm C. Jennings, Shanna B. Samels, Krisha S. Shah, Joya Maser, Medha Gupta, Jessica R. Barson

## Abstract

**Background:** Pituitary adenylate cyclase-activating polypeptide (PACAP) has been found to be involved in a wide range of motivated and affective behaviors. While the PACAP-38 isoform is more densely expressed than PACAP-27 in most of the brain, PACAP-27 is more highly expressed in the rodent paraventricular nucleus of the thalamus (PVT), where females also have greater expression than males. Notably, the role of PACAP-27 expression in cells of the PVT has not been explored.

**Methods:** Adult, female Long-Evans rats were injected in the PVT with an AAV to increase expression of PACAP or a control AAV. They were then investigated for subsequent gene and peptide levels of PACAP in the PVT; ethanol drinking and preference; sucrose drinking and preference; or locomotor activity in a novel chamber, behavior in a light-dark box, behavior in a novelty suppression of feeding test, locomotor activity in a familiar activity chamber, and behavior in a forced swim test.

**Results:** Gene expression of PACAP was significantly increased in the PVT by four weeks after injection with the PACAP AAV, and this resulted in a specific increase in levels of PACAP-27. Rats injected with the PACAP AAV demonstrated reduced drinking and preference for ethanol under the intermittent-access procedure compared to those injected with the control AAV. In contrast, rats injected with the PACAP AAV showed no significant difference in drinking or preference for sucrose, or in any affective behavior tested, except that they spent less time swimming in the forced swim test.

**Conclusions:** In light of the low overall level of expression of PACAP-27 in the brain, the ability of PACAP-27 in the PVT to control ethanol drinking, with minimal effects on other motivated or affective behaviors, supports the idea that compounds related to PACAP-27 should be investigated as potential therapeutics for the treatment of alcohol use disorder.

---

Pituitary adenylate cyclase-activating polypeptide (PACAP) is a pleiotropic peptide involved in phenomena that range from circadian rhythm, inflammation, and nociception to affective behavior, feeding, and intake of abused drugs (Harmar et al., 2012, Gargiulo et al., 2020). Largely, PACAP has been found to promote aversion, with central injection of PACAP significantly inhibiting food intake (Morley et al., 1992), and global knockout of PACAP at times being found to increase distance traveled in a novel environment (Hashimoto et al., 2001), reduce time spent in the dark chamber of a light-dark box (Gaszner et al., 2012), and decrease immobility in a forced swim test (Hattori et al., 2012). A connection between PACAP and alcohol intake has also been established. In social drinkers, a single-nucleotide polymorphism of the PACAP gene (*ADCYAP1*) has been associated with alcohol consumption (Kovanen et al., 2010), and in young women with problematic alcohol use, a single-nucleotide polymorphism of the PACAP receptor gene (*ADCYAP1R1*), has been associated with level of drinking behavior (Dragan et al., 2017). Similarly, in mice, global knockout of the PACAP gene has been found to result in increased intake of and preference for *ad libitum* ethanol (Tanaka et al., 2010). The brain areas through which PACAP acts to induce these changes in behavior are now being identified.

A number of regions in the limbic system have been found to express PACAP and / or its receptor and appear to mediate the effects of PACAP on alcohol drinking. With microinjections of PACAP and viral mediated knockdown of the PACAP receptor, these areas include the nucleus accumbens shell (Minnig et al., 2021, Gargiulo et al., 2021), the nucleus accumbens core (Minnig et al., 2022), and the bed nucleus of the stria terminalis (Ferragud et al., 2021). They also include the paraventricular nucleus of the thalamus (PVT). While the PACAP-38 isoform represents the large majority of PACAP in the brain (Arimura et al., 1991), it is PACAP-27 that is highly expressed in the rodent PVT (Gupta et al., 2018, Curtis et al., 2023). This is notable given the stronger connections made between aversion and PACAP-38 (Gargiulo et al., 2020). Expression of PACAP-27 in the rat PVT follows a gradient, with more PACAP-27^+^ cells being found in the posterior half of this nucleus (Gupta et al., 2018, Curtis et al., 2023). This peptide is also more highly expressed in the PVT of female rats and mice compared to males (Curtis et al., 2023). In male rats, levels of PACAP-27 in individual cells of the PVT have been found to be elevated at the end of an ethanol binge under the intermittent-access two-bottle-choice procedure (Gupta et al., 2018). While this suggests that PACAP in the PVT could function as a negative feedback signal to inhibit ethanol drinking, this remains to be determined.

The purpose of this study was to determine if PACAP in cells of the PVT could affect ethanol drinking and, if so, if it also influenced other motivated and affective behaviors. In female rats, which have higher baseline expression of PVT PACAP, an adeno-associated virus (AAV) was injected into the posterior PVT to increase expression of PACAP, and subsequent ethanol drinking, sucrose drinking, and affective behavior was then examined. We hypothesized that elevated levels of PACAP in cells of the PVT would inhibit ethanol drinking, but that it may have only minimal effects on sucrose drinking and affective behavior.

## MATERIALS AND METHODS

### Subjects

Adult, female Long-Evans rats (*N* = 83, 7 weeks on arrival at the facility, Charles River Laboratories International, Inc., Malvern, PA, USA) were individually housed in an AAALAC-accredited facility, on a 12-hour reversed light/dark cycle (lights off at 0900 h). They were given one week to acclimate to the facility. All animals received *ad libitum* chow (Laboratory Rodent Diet 5001, Lab Diet, St. Louis, MO, USA) and water throughout the study. Estrous cycle was not assessed, to minimize stress and reduce the chances of pseudo-pregnancy (Singletary et al., 2005, Lovick and Zangrossi, 2021). Experiments were approved by the Institutional Animal Care and Use Committee of Drexel University College of Medicine and followed the NIH Guide for the Care and Use of Laboratory Animals.

### Viruses

Viruses were designed and purchased from Vector Biolabs (Malvern, PA, USA). The AAV AAV9-CamKII-r-ADCYAP1-IRES-eGFP-WPRE (PACAP AAV) at a titer of about 1^e13^ gc/ml was used to increase expression of PACAP. The control AAV was AAV9-CamKII-eGFP-WPRE.

### Surgery

Rats were initially anesthetized with 5% isoflurane in 2 L/min oxygen and were then maintained under continuous anesthesia with 2 − 3% isoflurane in 1 L/min oxygen. Throughout the surgery, rats were kept on a hot water recirculating pad, to prevent hypothermia. Warm saline (5 ml s.c., Baxter Health Care Corp, Deerfield, IL, USA) was injected to prevent dehydration, and bupivicaine (2 mg/kg s.c., Hospira Worldwide, Lake Forest, IL, USA) was injected into the scalp prior to incision. Rats were injected into the posterior half of the PVT (2.8 mm posterior to Bregma, ±0.0 mm lateral to midline, 5.5 mm ventral to the level skull), with the PACAP AAV or control AAV, by lowering a 1 µL syringe (Hamilton Company, Reno, NV, USA) to the target depth, using a Kopf Model 5000 Microinjection Unit (Tujunga, CA, USA), and injecting 0.5 µL of the virus over 5 minutes, with an additional 5 minutes allowed for diffusion. The scalp incision was closed using 9 mm stainless steel wound clips (MikRon Precision Inc., Gardena, CA), and the skin was treated with Triple Antibiotic Ointment (MedPride, Bayonne, NJ, USA) and Lidocaine Cream (Curist, New York, NY, USA). Buprenorphine hydrocholoride (0.03 mg/kg s.c., Reckitt & Colman Inc, Slough, UK) was administered for post-operative analgesia. The wound clips were removed 7 - 10 days following surgery, under isoflurane anesthesia.

### Ethanol or Sucrose Drinking

Under the intermittent-access two-bottle-choice procedure adapted from Wise (1973) and Simms (2008), rats were given access to unsweetened 20% v/v ethanol or 2.5% w/v sucrose during three 24-hour-sessions per week in addition to *ad libitum* water and chow as described (Pirino et al., 2022, Gargiulo et al., 2021, Barson et al., 2015). Ethanol or sucrose was given each Monday, Wednesday, and Friday, one-and-a-half hours after dark onset. Ethanol intake was calculated as: (weight ethanol solution consumed (g) * (density ethanol * 0.20)) / rat body weight (kg). Sucrose intake was calculated as: (weight sucrose solution consumed (g) * (density sucrose * 0.025)) / rat body weight (kg). Ethanol and sucrose preference was calculated as: volume ethanol or sucrose solution consumed (ml) / (volume ethanol or sucrose solution consumed (ml) + volume water consumed (ml)). Food intake was calculated for Thursdays and Fridays, by weighing the food hopper, at the same time as the bottles, on Wednesdays, Thursdays, and Fridays. Animals were weighed on Tuesdays and Fridays.

### Quantitative Real-Time PCR (qRT-PCR)

Rats were sacrificed during the first half of their dark cycle, and their posterior PVT was dissected out to examine mRNA levels of PACAP (preproPACAP) using qRT-PCR, as previously described (Gargiulo et al., 2022, Curtis et al., 2023). The PVT sample was dissected as an inverted isosceles triangle directly ventral to the dorsal third ventricle, approximately 0.8 mm wide at the base, from a target slice that was made from Bregma −2.5 mm to −3.5 mm (Paxinos and Watson, 2005). Total RNA yield showed A260/A280 ratios between 1.80 and 2.21, indicating high purity. Target gene expression was quantified using the relative quantification method (ΔΔC_T_). The primers, used at a 200 nM concentration, were cyclophilin-A (housekeeping gene) forward: 5′-GTGTTCTTCGACATCACGGCT-3′, reverse: 5′-CTGTCTTTGGAACTTTGTCTGCA-3′; and PACAP (target gene) forward: 5′-GCCTCTCTGGTTGTGATTCCA-3′, reverse: 5′-GGTCATTCGCGGCTAGGAA-3′.

### Immunofluorescent Histochemistry

During the first half of their dark cycle, rats were deeply anesthetized with EUTHASOL® Euthanasia Solution (1 ml, i.p.) and perfused transcardially with 200 ml of ice-cold 0.9% sodium chloride and .02% Heparin (Sagent Pharmaceuticals, Schaumburg, IL, USA) solution followed by 250 ml of 4% paraformaldehyde in 0.1 M phosphate buffer, pH 7.4. Brains were then removed, post-fixed in 4% paraformaldehyde for 24 hours at 4 °C, cryoprotected in 30% sucrose for 2 − 4 days at 4 °C, and then frozen and stored at −80 °C. Brains were sliced by cryostat into 30 μm sections, which were stored at −20 °C in antifreeze solution (37.5% ethylene glycol, 20% sucrose in 0.03 M phosphate buffered saline (PBS)). Every sixth section through the posterior PVT was collected for processing and analysis, resulting in approximately 6 representative samples per brain.

To label PACAP, free-floating sections were rinsed for 10 min in 0.1% hydrogen peroxide to remove endogenous peroxidase activity. After rinsing in 0.1 M PBS, they were blocked for 90 min in 5% normal goat serum containing 0.5% Triton X-100 in PBS and incubated overnight at 4 °C in polyclonal guinea pig anti-PACAP-38 (1:200, OriGene Technologies, Inc, Rockville, MD, USA; cat# TA364386, lot# 022060) and polyclonal rabbit anti-PACAP-27 (1:200, LifeSpan BioSciences, Inc, Shirley, MA, USA; cat# LS-C144092, lot# 219923) primary antibodies. After the antibody incubation, the slices were rinsed in PBS and incubated for 2 h in goat anti-guinea pig (Alexa Fluor® 555) (1:200, Abcam, Cambridge, MA, USA; cat# ab150186, lot# GR128511-1) and goat anti-rabbit (Alexa Fluor® 647) (1:200, Abcam, Cambridge, MA, USA; cat# ab150079, lot# GR3444080-1) secondary antibodies in block solution. Alternate sections did not show immunofluorescence when processed with the antibodies omitted. Sections were mounted on slides and dried overnight in the dark, coverslipped with ProLong® Diamond Antifade Mountant with diamidino-2-phenylindole (DAPI; Life Technologies, Carlsbad, CA, USA), and allowed to set for at least 24 hours before imaging.

Imaging was conducted with a Leica DM5500 automated microscope (Buffalo Grove, IL, USA), and the images were captured with an Olympus DP71 high resolution digital color camera (Waltham, MA, USA) with Slidebook V6 image acquisition and analysis software (3i, Denver, CO, USA). Exposure conditions were the same for each slice processed with each primary antibody. Analyses were confirmed with a Leica SP8 VIS/405 HyVolution confocal microscope (Buffalo Grove, IL, USA). Quantification was conducted by an evaluator blind to the experimental condition of the subject. Evaluation of integrated density (area * mean gray value) for PACAP-27, PACAP-38, and AAV was performed with NIH software, ImageJ version 1.48v for Windows (Schneider et al., 2012). Each image in the appropriate channel was converted to grayscale, the PVT was isolated, and the threshold was adjusted, with the same limits used for each slice. Integrated density evaluates all labeling, including both cell bodies and fibers. Because of the variability in AAV labeling between subjects and across tissue slices, which did not follow a predictable pattern, data were evaluated as integrated density of each PACAP isoform as a function of integrated density of AAV labeling.

### Behavioral Testing

Unless otherwise noted, all behavioral testing was conducted in a sound- and light-attenuated room (< 5 lux), during the first half of the dark cycle. All rats were acclimated to the testing room for 5 minutes prior to each test.

#### Locomotor Activity

Locomotor activity was assessed in an automated activity chamber with an area of 43.2 cm x 43.2 cm and 42 cm high walls (Med Associates, Inc., St. Albans, VT, USA). Animals were placed at the center of the chamber and allowed to explore for 15 minutes, while ambulatory time and distance and vertical time (rearing) were measured via infrared beams (Pirino et al., 2020, Barson et al., 2015).

#### Light-Dark Box

A light/dark insert (Med Associates, Inc., St. Albans, VT, USA) was placed in the same chamber used for locomotor activity testing, to create a two-chamber light-dark box. A lamp was placed directly above the light chamber, creating a luminous intensity of approximately 400 lux in that chamber. Animals were placed in the light chamber and allowed to explore the light-dark box for 5 minutes. Time spent in the light chamber and number of entries into the two chambers were measured via infrared beams (Gargiulo et al., 2021, Pirino et al., 2022, Pirino et al., 2020). The first 5-minute period in the light-dark box was used for acclimation to the chamber, as we and others have found that behavior during the first time in the light-dark box is different from behavior during subsequent times (Rodgers and Shepherd, 1993, Pirino et al., 2022, Bouwknecht et al., 2004). A second 5-minute period in the light-dark box, given on the subsequent day, was used for the test.

#### Novelty Suppression of Feeding

Rats were acclimated to the novel diet (M&Ms^©^, Mars Inc, McLean, VA, USA) by being given one non-red-color M&M per day in the home cage, over three sequential days. Then, in the same chamber used for locomotor activity testing, three M&Ms^©^ were placed in one quadrant of the chamber, and *ad libitum* fed rats were placed in the opposite quadrant and allowed to explore the chamber for 5 minutes. Latency to enter the quadrant was measured via infrared beams, latency to first eating bout was recorded in real time by two evaluators blind to the experimental condition of the subject, and total food consumed was measured by weighing the novel food before and after each test.

#### Forced Swim Test

In a room at 35 lux, rats were placed in a cylinder that was 49 cm tall, with a 29 cm diameter, that contained 30 cm of water at 25 ± 2°C, and they were allowed to swim for 10 minutes, with 5 minutes for acclimation followed immediately by 5 minutes for testing. Time spent immobile (least amount of movement necessary to keep afloat), swimming (moving in the cylinder using hindlimbs), and climbing (thigmotactic behavior using forelimbs on the wall of the cylinder) were quantified from recorded videos by two evaluators blind to the experimental condition of the subject.

### Experimental Protocols

#### Experiment 1

To validate that the PACAP AAV increased levels of PACAP in the PVT, one set of rats was examined with qRT-PCR after being injected with the PACAP AAV or control AAV and being sacrificed at 1 day or 28 days after surgery, or they did not undergo surgery and were left as cage controls (*N* = 30, *n* = 6/group). A second set of rats (*N* = 15, *n* = 7 PACAP AAV, 8 control AAV) was examined with immunohistochemistry after being injected with the PACAP or control AAV, tested for affective behavior, and sacrificed at 40 days after surgery.

#### Experiment 2

To determine if the PACAP AAV could affect ethanol drinking, rats were injected with the PACAP or control AAV (*N* = 18, *n* = 9/group) and, starting 21 days after surgery, were given ethanol for 8 weeks under the intermittent-access two-bottle-choice paradigm. Data from 3 rats (2 PACAP AAV, 1 control AAV) were not included in the analysis, due to injections having been > 0.5 mm from the target region. To determine if the effects on ethanol were substance-specific, a second set of rats was injected with the PACAP or control AAV (*N* = 17, *n* = 9 PACAP AAV, 8 control AAV) and, starting 21 days after surgery, given sucrose for 8 weeks under the intermittent-access two-bottle-choice paradigm. At the end of their time drinking, rats were sacrificed, and their injection placement was verified using 30 μm brain sections under a Leica DM5500 microscope.

#### Experiment 3

To determine if the PACAP AAV could impact affective behavior, the rats that were ultimately used for immunohistochemistry in Experiment 1 (*N* = 18, *n* = 9/group) were tested starting 21 days after surgery in the following order: locomotor activity in a novel chamber (day 21), light-dark box (acclimation on day 22, test on day 25), novelty suppression of feeding test (acclimation on days 25 – 27, test on day 28), locomotor activity (day 32 or 33), and forced swim test (day 36). Injection placement was verified on 30 μm brain sections under a Leica DM5500 microscope, using alternate sections from those used for immunohistochemistry.

### Data Analysis

Sphericity of the data was determined using Mauchly’s test, and a Greenhouse-Geisser correction was used when sphericity was violated. Data from Experiment 1 were analyzed using a one-way ANOVA (qRT-PCR) or mixed ANOVA (immunofluorescent histochemistry), with group as the between-subject factor and PACAP isoform as the within-subject factor. Data from Experiment 2 were analyzed using a mixed ANOVA, with group as the between-subject factor and day as the within-subject factor. For ANOVAs, significant main and interaction effects were followed up with Sidak pairwise comparison tests. Data from Experiment 3 were analyzed using independent-samples two-tailed *t*-tests. For data that were manually obtained from two evaluators, the average value was used for analysis. All data were analyzed using SPSS (Version 29, IBM, Armonk, NY, USA) and all graphical figures were made using GraphPad Prism (Version 10, Boston, MA, USA). Significance was determined at *p* < 0.05 and all *p* values are reported with up to three significant figures. Data are reported as mean ± standard error of the mean (S.E.M.).

## RESULTS

### Experiment 1: Verification of PVT PACAP overexpression

To validate that the PACAP AAV increased levels of PACAP in the PVT, rats were injected with the PACAP AAV or control AAV and were sacrificed at 1 day or 28 days after surgery, or they were left as cage controls (*N* = 30, *n* = 6/group), and PACAP mRNA in the PVT was examined with qRT-PCR. A one-way ANOVA revealed that there was a significant difference between groups [*F*(4, 30) = 4.46, *p* = 0.007], and pairwise comparisons demonstrated that this was due to significantly higher levels of PACAP mRNA in the PACAP AAV group that was sacrificed at 28 days, compared to all other groups (*p* = 0.016 – 0.037) (**Figure 1A**). There were no significant differences between any other groups.

**Fig 1.**
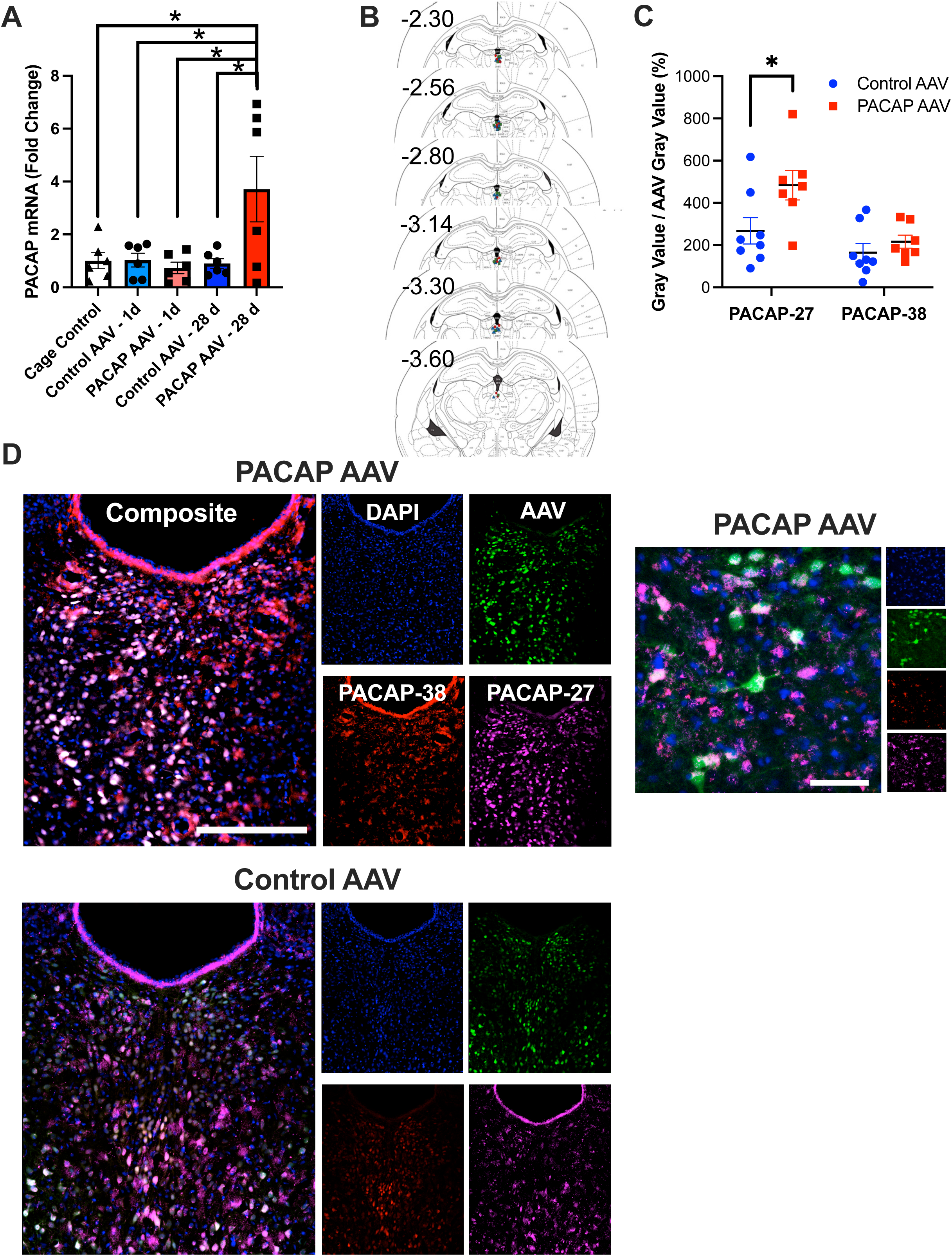
Expression of PACAP in the PVT after injection with the PACAP AAV (Experiment 1). **A.** Rats injected with the PACAP AAV showed significantly higher levels of PACAP mRNA in the PVT by 28 days after injection, compared to rats injected with the control AAV, those examined at 1 day after injection, and those left as cage controls (*N* = 30, *n* = 6/group). **B.** Histological examination revealed that injections were made into the middle or posterior subregion of the PVT. Blue triangles = Experiment 1 / 3 (immunofluorescent histochemistry / affective behavior), green squares = Experiment 2 (ethanol drinking), red circles = Experiment 2 (sucrose drinking). **C.** Rats injected with the PACAP AAV showed significantly higher levels of PACAP-27 but not PACAP-38 as a function of AAV labeling by 40 days after injection, compared to rats injected with the control AAV (*N* = 15, *n* = 7 PACAP AAV, 8 control AAV). **D.** Confocal photomicrographs showing 4’,6-diamidino-2-phenylindole (DAPI, blue), the AAV (green), PACAP-38 (red), and PACAP-27 (fuschia) in the posterior PVT of a rat injected with the PACAP AAV (top left) and a rat injected with the control AAV (bottom left), as well as a high magnification image of a segment of the tissue slice from the rat injected with the PACAP AAV (top right). Scale bar in main image = 250 μm. Scale bar in high magnification image = 50 μm. Data are mean ± S.E.M., \**p* < 0.05.

To determine if a specific isoform of PACAP peptide was increased in the PVT by the PACAP AAV, rats were injected with the PACAP AAV or control AAV and were sacrificed at 40 days after surgery, following behavioral testing (*N* = 15, *n* = 7 PACAP AAV, 8 control AAV). Histological examination revealed that injections were made into the middle or posterior subregion of the PVT, between −2.30 and −3.60 mm relative to Bregma (**Figure 1B**). The AAVs appeared to have a radial spread of ∼0.6 mm, such that they primarily transduced cells in the posterior half of the PVT. A mixed ANOVA revealed that there was a significant difference in the overall labeling of PACAP-27 versus PACAP-38, as a function of AAV labeling [*F*(1, 13) = 31.47, *p* < 0.001], with a significant interaction between isoform labeling and AAV group [*F*(1, 13) = 6.16, *p* = 0.028] (**Figure 1C, D**). Pairwise comparisons demonstrated that this was due to significantly higher labeling of PACAP-27 than PACAP-38 in both the control AAV (*p* = 0.039) and the PACAP AAV (*p* < 0.001) groups. Moreover, while the difference in overall PACAP labeling between the control and PACAP AAV groups did not attain statistical significance [*F*(1, 13) = 3.77, *p* = 0.074], there was significantly higher labeling of PACAP-27 (+80%, *p* = 0.038) and not PACAP-38 (+31%, *p* = 0.355) in the PVT compared to the control AAV group. These results together demonstrate that injection of the PACAP AAV into the rat PVT results not just in increased levels of PACAP mRNA, but a specific increase in levels of PACAP-27 in the PVT.

### Experiment 2: Effects of PVT PACAP overexpression on ethanol and sucrose drinking

To determine if the PACAP AAV could affect ethanol drinking, rats were injected with the PACAP AAV or control AAV (*N* = 15, *n* = 7 PACAP AAV, 8 control AAV) and were given ethanol for 8 weeks under the intermittent-access two-bottle-choice procedure. For ethanol drinking, a mixed ANOVA showed no significant difference between groups [*F*(1, 13) = 3.40, *p* = 0.088], but a significant difference between weeks [*F*(2.99, 38.80) = 5.941, *p* = 0.002], and a significant interaction between group and week [*F*(7, 91) = 2.43, *p* = 0.025]. Pairwise comparisons revealed that, whereas rats injected with the control AAV drank less on Week 1 than on Weeks 3 (*p* = 0.002), 5 (*p* = 0.002), 6 (*p* = 0.001), 7 (*p* = 0.003), and 8 (*p* = 0.008), those injected with the PACAP AAV showed no significant difference in drinking across weeks (*p* = 0.873 – 1.000). As a result, rats injected with the PACAP AAV drank less ethanol than those injected with the control AAV on Week 6 (*p* = 0.032) and Week 8 (*p* = 0.044) (**Figure 2A**). Similarly, for ethanol preference, there was a trend for a significant difference between groups [*F*(1, 13) = 4.48, *p* = 0.054], a significant difference between weeks [*F*(3.24, 42.16) = 7.87, *p* < 0.001], and a significant interaction between group and week [*F*(7, 91) = 3.42, *p* = 0.003] (**Figure 2B**). Pairwise comparisons revealed that rats injected with the control AAV showed a lower ethanol preference on Week 1 than on Weeks 3 – 8 (*p* = 0.045 - *p* < 0.001) while those injected with the PACAP AAV showed no significant difference in preference across weeks (*p* = 0.551 – 1.000). Rats injected with the PACAP AAV showed a lower ethanol preference than the control AAV group on Week 6 (*p* = 0.045), Week 7 (*p* = 0.020), and Week 8 (*p* = 0.006). In contrast to ethanol drinking and preference, food intake was not significantly different between groups [*F*(1, 12) = 0.62, *p* = 0.447] or weeks [*F*(2.47, 29.60) = 1.13, *p* = 0.351] (**Figure 2C**). Body weight was also not significantly different between groups [*F*(1, 13) < 0.01, *p* = 0.966] and, although there was a significant difference between weeks [*F*(1.64, 21.35) = 149.43, *p* < 0.001], there was no significant interaction between groups and weeks [*F*(7, 91) = 0.71, *p* = 0.666] (**Figure 2D**).

**Fig 2.**
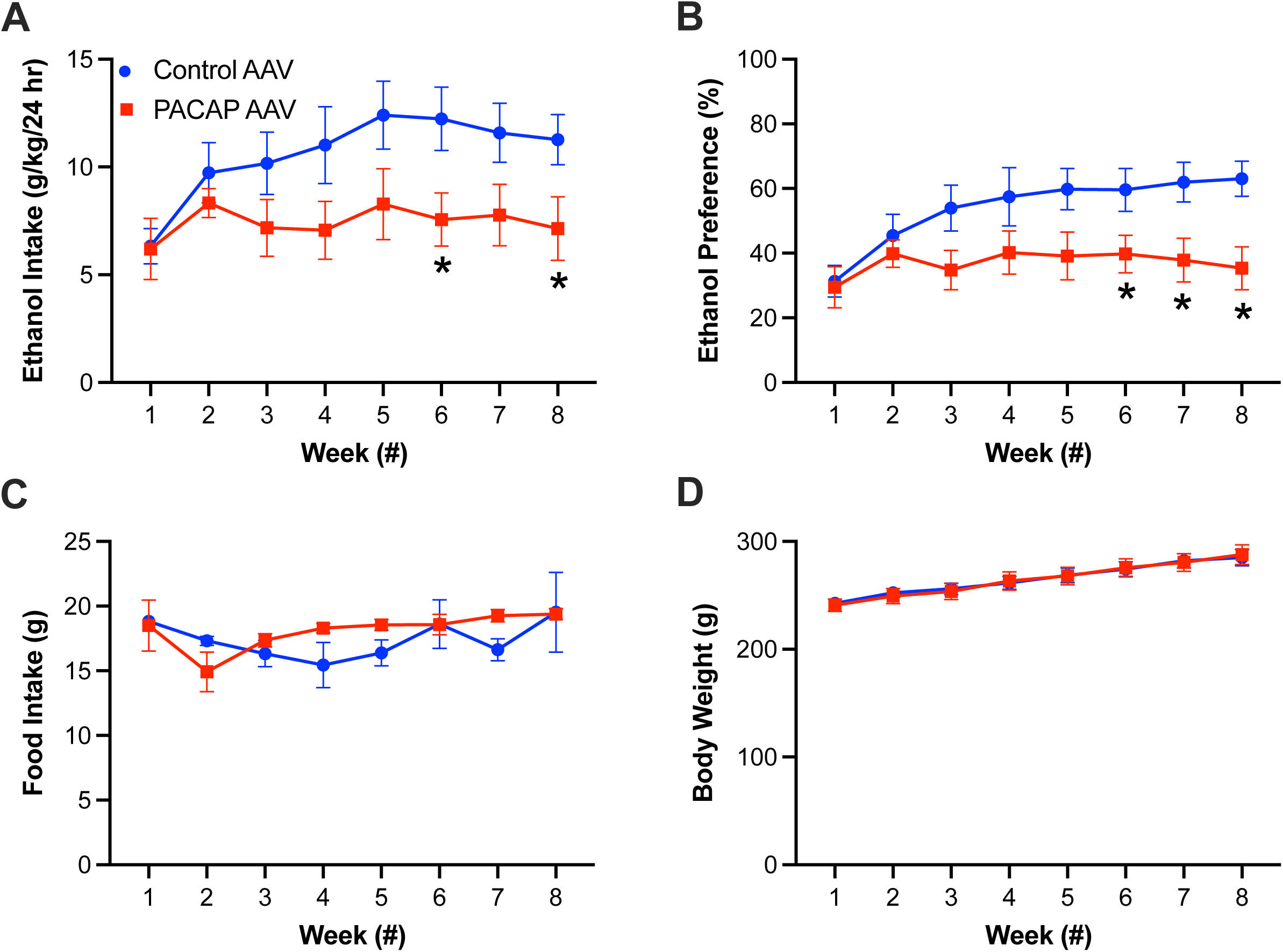
Ethanol drinking in rats injected with a PACAP AAV or control AAV (*N* = 15, *n* = 7 PACAP AAV, 8 control AAV) (Experiment 2). **A.** Rats injected with a PACAP AAV drank significantly less ethanol than rats injected with a control AAV during drinking Weeks 6 and 8. **B.** Rats injected with the PACAP AAV showed a lower ethanol preference than rats injected with a control AAV during drinking Weeks 6, 7, and 8. **C.** Rats injected with a PACAP AAV showed no significant difference in chow intake than rats injected with a control AAV during the 8 weeks of ethanol access. **D.** Rats injected with a PACAP AAV showed no significant difference in body weight than rats injected with a control AAV during the 8 weeks of ethanol access. Data are mean ± S.E.M., \**p* < 0.05 vs. control AAV group.

To determine if the effects observed on ethanol were substance-specific, other rats were injected with the PACAP AAV or control AAV (*N* = 17, *n* = 9 PACAP AAV, 8 control AAV) and given sucrose for 8 weeks. For sucrose drinking, a mixed ANOVA showed no significant difference between groups [*F*(1, 14) = 0.05, *p* = 0.829] and, although there was a significant difference between weeks [*F*(7, 98) = 7.61, *p* < 0.001], there was no significant interaction between groups and weeks [*F*(7, 98) = 0.85, *p* = 0.551] (**Figure 3A**). Similarly, for sucrose preference, there was no significant difference between groups [*F*(1, 15) = 0.39, *p* = 0.541], a significant difference between weeks [*F*(3.32, 49.78) = 7.67, *p* < 0.001], and no significant interaction between groups and weeks [*F*(7, 105) = 0.69, *p* = 0.684] (**Figure 3B**). As with ethanol drinkers, food intake in these sucrose drinkers was not significantly different between groups [*F*(1, 15) = 0.09, *p* = 0.767] and, although there was a significant difference between weeks [*F*(3.91, 58.61) = 4.79, *p* < 0.001], the interaction effect showed only a trend for significance [*F*(7, 105) = 2.03, *p* = 0.058] and pairwise comparisons revealed no significant difference between the PACAP AAV group and the control AAV group on any single week (*p* = 0.198 – 0.979) (**Figure 3C**). Body weight was also not significantly different between groups [*F*(1, 15) = 1.15, *p* = 0.300] and, although there was a significant difference between weeks [*F*(2.14, 32.16) = 42.15, *p* < 0.001], there was no significant interaction effect between groups and weeks [*F*(7, 105) = 1.34, *p* = 0.239] (**Figure 3D**). These results demonstrate that overexpression of PACAP in the PVT can lead to a reduction in ethanol intake and preference, with no effect on sucrose or chow intake or on body weight.

**Fig 3.**
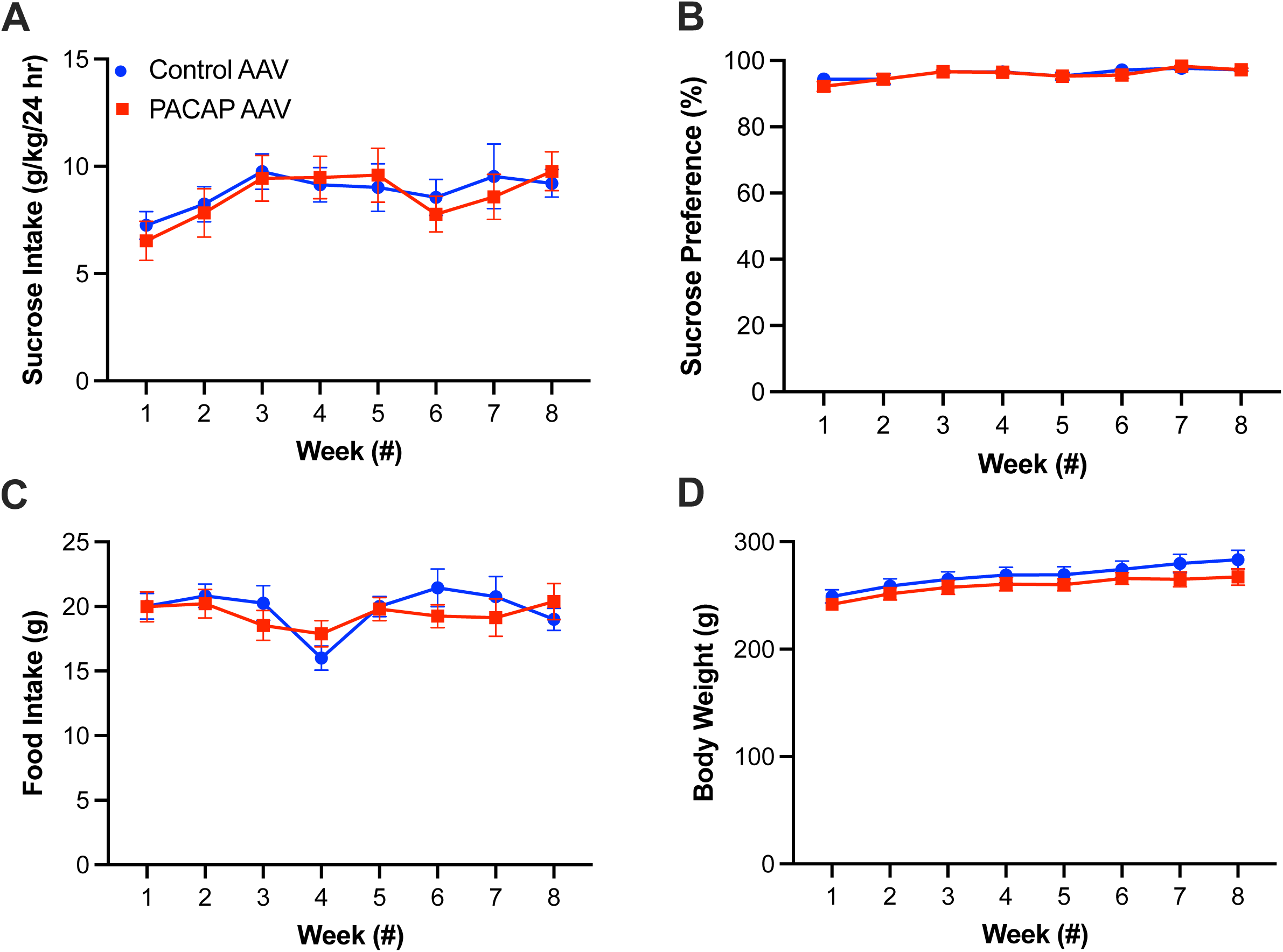
Sucrose drinking in rats injected with a PACAP AAV or control AAV (*N* = 17, *n* = 9 PACAP AAV, 8 control AAV) (Experiment 2). **A.** Rats injected with a PACAP AAV drank sucrose at levels that were not significantly different than rats injected with a control AAV. **B.** Rats injected with the PACAP AAV had sucrose preference levels that were not significantly different than rats injected with a control AAV. **C.** Rats injected with a PACAP AAV showed no significant difference in chow intake than rats injected with a control AAV during the 8 weeks of sucrose access. **D.** Rats injected with a PACAP AAV showed no significant difference in body weight than rats injected with a control AAV during the 8 weeks of sucrose access. Data are mean ± S.E.M.

### Experiment 3: Effects of PVT PACAP overexpression on affective behavior

To determine if the PACAP AAV could impact affective behavior, rats were injected with the PACAP AAV or control AAV (*N* = 18, *n* = 9/group) and, starting 21 days after surgery, were run through a battery of behavioral tests. For locomotor activity in a novel activity chamber, there was no significant difference between groups for ambulatory time [*t*(16) = 0.55, *p* = 0.593], ambulatory distance [*t*(16) = 0.70, *p* = 0.493], or vertical time [*t*(15) = 0.42, *p* = 0.683] (**Figure 4A**). For behavior in a light-dark box, there was no significant difference between groups for time spent in the light chamber [*t*(16) = −0.67, *p* = 0.511] or entries into the light chamber [*t*(16) = −0.92, *p* = 0.371] (**Figure 4B**). For behavior in a novelty suppression of feeding test, there was no significant difference between groups for total food intake [*t*(10.00) = −1.92, *p* = 0.084], latency to enter the quadrant with the food [*t*(16) = −0.80, *p* = 0.435], or latency to start eating [*t*(15) = 1.35, *p* = 0.196] (**Figure 4C**). Similarly, for locomotor activity in a familiar activity chamber, there was no significant difference between groups for ambulatory time [*t*(16) = −0.64, *p* = 0.533], ambulatory distance [*t*(16) = −0.53, *p* = 0.601], or vertical time [*t*(15) = −0.81, *p* = 0.431] (**Figure 4D**). In contrast, for behavior in a forced swim test, while there was no significant difference between groups for time spent immobile [*t*(16) = −2.01, *p* = 0.062] or time spent climbing [*t*(16) = −0.24, *p* = 0.817], rats injected with the PACAP AAV spent significantly less time swimming than rats injected with the control AAV [*t*(11.16) = −2.24, *p* = 0.046] (**Figure 4E**). All together, these data suggest that overexpression of PACAP in the PVT leads to very few significant changes in affective behavior, although it does alter behavior in a forced swim test.

**Fig 4.**
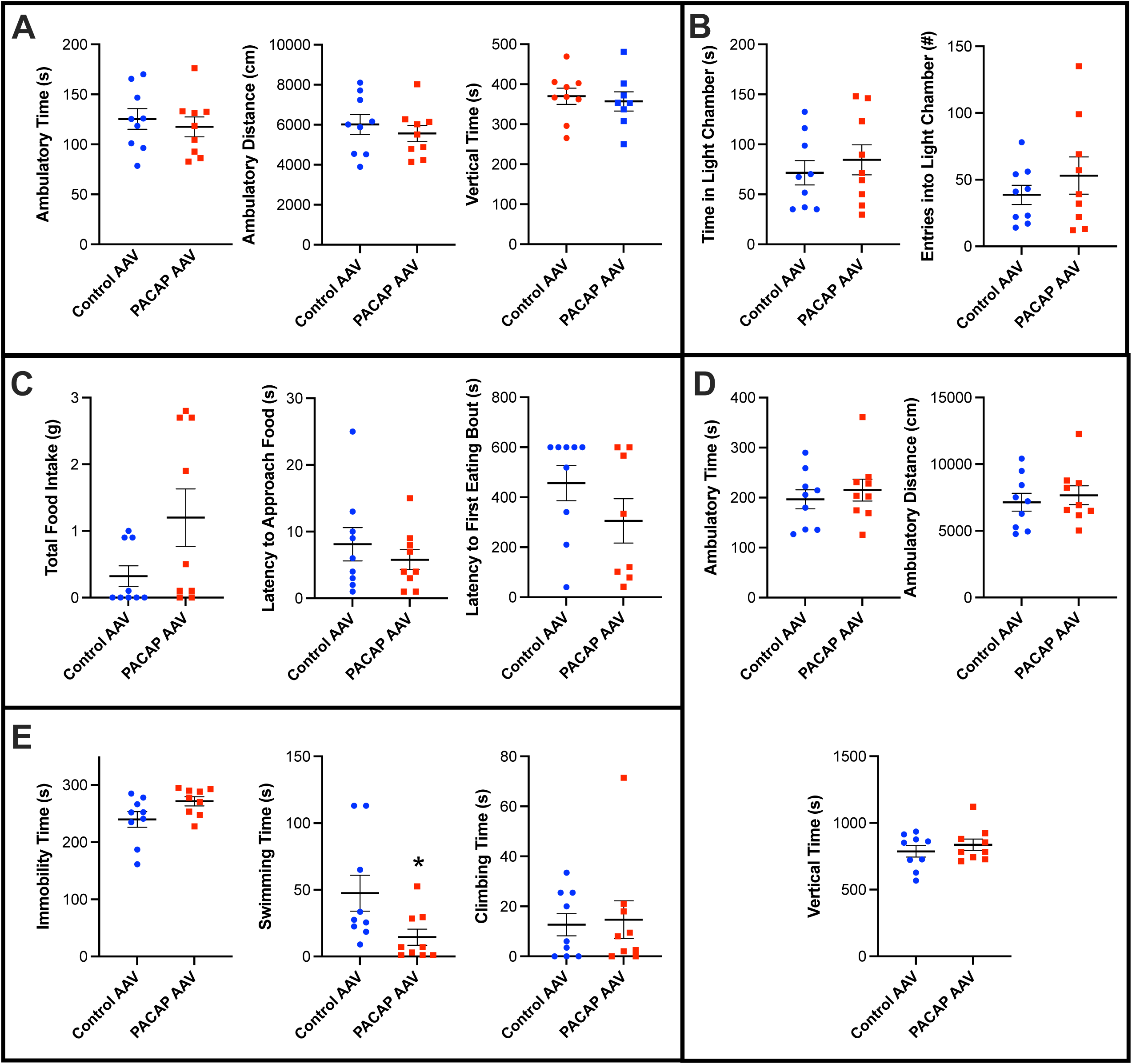
Affective behavior in rats injected with a PACAP AAV or control AAV (*N* = 18, *n* = 9/group) (Experiment 3). **A.** Rats injected with a PACAP AAV showed no significant difference in behavior in locomotor activity in a novel activity chamber compared to rats injected with a control AAV. **B.** Rats injected with a PACAP AAV showed no significant difference in behavior in a light-dark box compared to rats injected with a control AAV. **C.** Rats injected with a PACAP AAV showed no significant difference in behavior in a novelty suppression of feeding test compared to rats injected with a control AAV. **D.** Rats injected with a PACAP AAV showed no significant difference in behavior in locomotor activity in a familiar activity chamber compared to rats injected with a control AAV. **E.** Rats injected with a PACAP AAV showed significantly less time swimming in a forced swim test than rats injected with a control AAV. Data are mean ± S.E.M., \**p* < 0.05 vs. control AAV group.

## DISCUSSION

In this study, we used an AAV that, after injection into the middle/posterior PVT of female rats, elevated local gene expression of PACAP, and specifically increased peptide levels of PACAP-27 in the PVT. Compared to rats injected with a control AAV, this PACAP AAV led to reduced ethanol drinking and preference for 20% ethanol under the intermittent-access two-bottle-choice procedure. It did not alter sucrose drinking or preference, under a similar intermittent-access procedure. The PACAP AAV also did not lead to changes in behavior in tests of locomotor activity or approach-avoidance behavior, but it did alter swimming behavior in a forced swim test. These results together suggest that PACAP in the PVT reduces ethanol drinking in a manner that is relatively behaviorally specific.

The finding that the PACAP AAV specifically increased levels of PACAP-27 in the PVT is consistent with the known expression of prohormone processing enzymes in the PVT. The PACAP precursor (preproPACAP) can be independently processed into PACAP-27 or PACAP-38 by the prohormone processing enzymes SPC2 (also called PC2 and RPC2) or SPC3 (also called PC1, PC3, and BPD), respectively (Li et al., 1999). In the PVT of the adult rat, it is SPC2 and not SPC3 that shows abundant expression (Birch et al., 1994). That is, the enrichment of SPC2 in cells of the PVT allowed for specific processing of PACAP mRNA into PACAP-27. In line with this, using validated isoform-specific antibodies, we have previously found in both male and female rats that PACAP-27 is the predominant PACAP isoform in the PVT, with 44 – 52% of all PVT cells expressing PACAP-27, in contrast to 10 – 17% of cells that express PACAP-38 (Gupta et al., 2018, Curtis et al., 2023). While we cannot rule out that the behavioral effects observed in this study are due to increased levels of PACAP-38 in the PVT, it appears likely that they are largely due to PACAP-27.

Rats with elevated levels of PACAP in the PVT as a result of injection with the PACAP AAV showed reduced ethanol intake and preference in later weeks of access to 20% ethanol under the intermittent-access two-bottle-choice procedure, compared to rats injected with the control AAV. Specifically, the difference between PACAP AAV- and control AAV-injected animals emerged at Week 6 and continued during Weeks 7 and 8 of the eight-week ethanol access. Notably, control rats showed an escalation of ethanol drinking and preference across weeks that was consistent with our prior observations of increased ethanol intake and preference across weeks under the intermittent-access procedure (Pirino et al., 2022, Pandey and Barson, 2020). Therefore, the PACAP AAV appears to have affected ethanol drinking by preventing the escalation of drinking. The reduced drinking from PACAP relative to a control is consistent with prior literature that used various ethanol access schedules and other means of manipulating the PACAP system. In male mice with *ad libitum* access to ethanol, global knockout of the PACAP gene lead to increased intake of and preference for ethanol (Tanaka et al., 2010); in male and female Long-Evans rats, microinjection of PACAP-27 into the nucleus accumbens shell reduced drinking under the intermittent access procedure (Gargiulo et al., 2021); and in alcohol-preferring male Scr:sP rats, knockdown of the PAC1 receptor in the nucleus accumbens shell led to increased ethanol drinking and preference under a self-administration paradigm (Minnig et al., 2021). It should be noted, however, that PACAP can also promote ethanol drinking, as has been observed in animals that approach dependence. For example, a PACAP-38 receptor antagonist reduced ethanol self-administration when injected into the bed nucleus of the stria terminalis of male Wistar rats after chronic intermittent access to ethanol vapor (Ferragud et al., 2021), and this same compound also reduced ethanol self-administration when injected into the nucleus accumbens core of high-drinking, alcohol-preferring male Scr:s rats (Minnig et al., 2022). The lack of effect of the PACAP AAV on sucrose drinking is also consistent with prior literature. For example, the same knockout of the PACAP gene in mice that increased preference for ethanol failed to increase preference for sucrose (Tanaka et al., 2010), and the same injection of PACAP-27 into the nucleus accumbens shell in rats that reduced ethanol drinking failed to significantly affect sucrose drinking (Gargiulo et al., 2021). All together, the present results demonstrate that an increase in levels of PACAP in cells of the PVT prevents an escalation of ethanol drinking under the intermittent access procedure, and it does so in a manner that is substance-specific.

In contrast to the effects on ethanol drinking, injection of a PACAP AAV in the PVT led to no significant effects on measures of locomotor activity or measures of approach-avoidance or anxiety-like behavior (activity in a novel chamber, behavior in light-dark box, feeding in a novelty suppression of feeding test) (Calhoon and Tye, 2015), but it did lead to reduced swimming in an inescapable situation (forced swim test) (Commons et al., 2017, Armario, 2021). Prior examination of effects of PACAP on locomotor activity and behavior in approach-avoidance tests have yielded inconsistent results, with intracerebroventricular injection of PACAP-38 and global knockout of the PACAP gene leading to reductions, increases, or no effects on distance traveled or activity counts in a familiar or novel enclosure (Masuo et al., 1995, Hattori et al., 2012, Marquez et al., 2009) and on time in the chambers of a light-dark box (Hattori et al., 2012, Gaszner et al., 2012). Similarly, chemogenetic activation of PACAP^+^ cells in the lateral habenula was not found to affect novelty-induced feeding suppression (Levinstein et al., 2022). Of note, we have found that injection of PACAP-38 but not PACAP-27 into the nucleus accumbens shell reduced time spent in the light chamber of a light-dark box (Gargiulo et al., 2021). Thus, although the PVT participates in avoidance or anxiety-like behavior (Barson et al., 2020), and the PVT sends major projections to the nucleus accumbens shell (Li et al., 2024, Li et al., 2021, Dong et al., 2017), the ability of the PVT to affect these behaviors may not occur through release of PACAP-27. On the other hand, while intracerebroventricular injection of PACAP-38 and global knockout of the PACAP gene have also led to mixed effects on immobility in a forced swim test (Tajiri et al., 2012, Seiglie et al., 2015, Hattori et al., 2012, Gaszner et al., 2012), prior research found that chemogenetic activation of PVT cells increased immobility in this test (Cui et al., 2024) while inhibition reduced it (Kato et al., 2019). In the present study, we found that an increase of PACAP in the PVT led to reduced time swimming, although it did not lead to a significant effect on immobility. While immobility in the forced swim test may reflect a passive coping strategy to an inescapable aversive situation, swimming may instead reflect an active strategy (Commons et al., 2017, Armario, 2021). Our results therefore suggest that PACAP in the PVT may counteract adaptive adjustment, rather than promote maladaptive strategies.

A major caveat of our results is that only females were tested in these studies. We targeted this population, given that females have higher baseline expression of PACAP in the PVT than males (Curtis et al., 2023), and they also drink more ethanol under the intermittent access procedure (Pirino et al., 2022, Morales et al., 2015, Li et al., 2019). Of note, this elevated drinking is not dependent on estrous cycle stage (Priddy et al., 2017, Li et al., 2019). In light of our findings that chronically increased levels of PACAP in the PVT prevent an escalation of ethanol drinking, we believe that baseline PACAP levels alone cannot be the determining factor of ethanol drinking. Indeed, ethanol intake itself appears to elevate levels of PACAP in the PVT (Gupta et al., 2018), and other chronic but not acute manipulations similarly elevate levels of PACAP in other brain regions (Lezak et al., 2014, Hannibal et al., 1995, Hannibal et al., 1999). Therefore, we hypothesize that there may be a dynamic interaction between ethanol drinking and levels of PACAP that determines level of ethanol drinking in a given bout.

In summary, this study reports that chronic elevation of PACAP levels in the PVT of female rats prevents an escalation of ethanol drinking and preference under the intermittent-access two-bottle-choice procedure and it also reduces active coping in a test of behavioral despair, but it leads to no significant changes in measures of sucrose drinking, locomotor activity, or approach-avoidance / anxiety-like behavior. In identifying the ability of PACAP-27 in the PVT to control ethanol drinking with minimal effects on other motivated or affective behaviors, these results lend support to the idea that compounds related to PACAP-27 should be investigated as potential therapeutics for the treatment of alcohol use disorder.

## ACKNOWLEDGMENTS

This research was supported by the National Institute on Alcohol Abuse and Alcoholism under Award Numbers R01AA028218 (J.R.B.), and F31AA031427 (B.A.C). The content is solely the responsibility of the authors and does not necessarily represent the official views of the NIH. The authors declare no conflict of interest.

## Notes

### Competing Interest Statement

The authors have declared no competing interest.

